# Association study of long-term kidney transplant rejection using whole-exome sequencing

**DOI:** 10.1101/444497

**Authors:** Louis Gioia, Sunil Kurian, Tony S. Mondala, Laia Bassaganyas, Pui-Yan Kwok, Daniel R. Salomon, Andrew I. Su

## Abstract

Long-term renal allograft rejection is the most common outcome in kidney transplantation. Continuing the crusade to extend allograft function after the first year post-transplantation, we attempted to associate genetic factors that might contribute to long-term allograft outcomes by sequencing the exomes of patients diagnosed with chronic allograft nephropathy/interstitial fibrosis and tubular atrophy. A variety of association analyses were employed, but these analyses failed to identify statistically significant associations. The study was underpowered to detect the association of rare genomic variants with small effect sizes. However, it confirmed previous reports of the absence of large effects from common variants. We have made both the study data and analysis workflow available for public use, and we hope that these resources will help to power future meta-analyses that may detect smaller effects.

## 1 Introduction

Renal transplantation is the best treatment option for most patients with end-stage renal disease. In 2017, nearly 20,000 kidney transplants were performed, representing over half of all organ transplant procedures (OPTN data as of September 5, 2018). Advancements in post-transplantation immunosuppressive therapies have greatly improved short-term allograft survival rates, with 1-year graft survival rates of over 90% for both living and deceased donors^1^. Now, as acute rejection and early graft loss have become less common, the current focus is on the prevention and treatment of longer-term complications and improving long-term graft outcomes.

The most common cause of graft loss after the first year post-transplant is chronic allograft nephropathy (CAN). CAN is defined as a general and still poorly understood clinical entity, characterized by progressive deterioration of organ function in the transplanted kidney^2^. CAN has been given various nomenclatures based on the clinical or histological definition of the underlying disease/pathology^3–6^, but for the sake of simplicity and consistency we will refer to the entity as CAN. While there has been an increased interest in CAN research, a clear mechanistic explanation for the clinical entity has not emerged. Rather, studies have indicated that CAN represents a spectrum of immune and nonimmune processes between the host and allograft, causing injury to the kidney in a progressive manner^7^.

Due to the inherent ambiguity in the definition of CAN, the clinical entity has been redefined in histological terms as interstitial fibrosis and tubular atrophy (IFTA) of unknown etiology^8^. Biopsy studies place the incidence of IFTA around 50% for transplants after 1 year, 70% after 2 years, and near 100% after 10 years^9^. These studies also reveal that IFTA severity correlates with allograft dysfunction and progresses over time. However, the progression of IFTA does not seem to proceed in a linear manner, supporting the idea that the processes driving IFTA are dynamic and exposing the need for further research into the mechanisms responsible for CAN/IFTA progression.

Many factors are known to influence long-term renal allograft function, and genetic factors have been especially suspected due to the success of kidney transplantation between identical twins^10,11^. However, to date, the only genetic variants that have been shown to be significantly associated with renal allograft dysfunction lie in the HLA region^12^. Despite decades of research, the role of the HLA region in transplant outcomes is still under debate. Though the effort taken to find compatible transplants over the last 25 years are commendable, the return on those efforts has been modest at most. The sheer numbers of HLA antigens and molecular subtypes, as well as their combinations and uneven distributions among racial and ethnic populations, makes matching difficult. This is particularly relevant for transplant centers, which must make efforts to be more inclusive of lower priority or extended criteria kidneys rather than discarding them. Continuing the search for genetic variants that contribute to extended kidney transplant outcomes, we sought to identify genetic variants that are associated with the specific CAN/IFTA phenotype through a case/control association study. For genetic variant identification, we employed whole-exome sequencing due to its balance between genome coverage and affordability^13^.

## 2 Materials and Methods

### 2.1 Study participant samples

260 peripheral blood samples were collected from patients who either displayed the extreme CAN/IFTA phenotype (> Banff grade I according to the Banff ’05 histological criteria) or had healthy transplants after 3 years^8^. Three cohorts of patients were selected, each of which contained near equal proportions of the two phenotypes. The first cohort consisted of 100 patients of varied self-reported ethnicity, the second consisted of 60 patients who self-identified as “Caucasian/European” (referred hereafter to as “white”), and the third consisted of 100 patients who self-identified as “African-American” (“black”). In total, 30 samples were removed from the analysis due to errors in sample metadata and/or issues with sequencing quality. Of the remaining 230 samples, 118 were CAN and 112 were excellent transplants (TX). All patient samples and clinical data were collected with approved Institutional Review Board (IRB) consents as part of the U19AI063603-01.

### 2.2 Sequencing and variant calling

Genomic DNA was extracted from whole blood using QIAGEN Puregene Blood Kits (Valencia, CA) according to the manufacturer’s protocol. Whole exome sequencing library preparation was performed using the KAPA^®^ Hyper Prep Kit (Pleasanton, CA). Exome capture was performed using the Nimblegen SeqCap EZ Human Exome Library v3.0 Kit (Madison, WI) according to the manufacturer’s protocols. Sequencing was performed on an Illumina HiSeq 2000 sequencer with a paired-end read length of 100 bp in the Genomics Core Facility at UCSF.

The sequencing reads from each sample were aligned to the hg19 human reference genome using the BWA-MEM algorithm from the Burrows-Wheeler Aligner (BWA) software package, version 0.7.10^14^. To prepare the alignment files for variant calling, read groups were assigned and duplicate reads were marked using the Picard Tools software from the Broad Institute (http://broadinstitute.github.io/picard/). Single-nucleotide variants and small insertions/deletions (indels) were identified using the Genome Analysis Toolkit (GATK) version 3.3 software, according to the Broad Institute’s best-practices workflow^15^. The Mills and 1000Genomes gold standard indels, the 1000Genomes Phase I indel calls, and the dbSNP build 137 data sets were used as known sites for indel realignment and base quality score recalibration. Joint variant calling was performed using the GATK HaplotypeCaller in GVCF mode, followed by joint genotyping to produce a group variant calling format (gVCF) file for each analysis cohort.

### 2.3 Quality control

In order to minimize labeling errors, the sex of each sample was determined using the Plink 1.9 software package^16^, and these sequence-based sex labels were compared to the original sex annotations. Sequence-based sexes were determined by computing a heterozygosity F-statistic estimate for variant calls on the X chromosome for each sample and visualizing the F-statistic distributions within each cohort to identify males and females. All sequencing cohorts displayed distinct and separate distributions for the male and female F-statistics. Samples with calculated sexes that differed from their original labels were then removed from the analysis—7 samples were removed from the first cohort, 8 samples were removed from the second cohort, and 3 samples were removed from the third sequencing cohort.

### 2.4 Association analyses

The Plink 1.9 software was used to run a Fisher’s Exact Test with Lancaster’s mid-p adjustment for each variant^16^. Quantile-quantile and Manhattan plots were created with the “qqman” R package^17^. An adaptive Monte Carlo permutation was also performed for each variant site, producing high-confidence permuted p-values.

Associations at the gene level were tested using a sequence kernel association test for the combined effect of rare and common variants (SKAT_CommonRare) for each analysis cohort^18^. SNP sets with gene annotations for each GATK-called variant were created using the hg19 gene range list provided by Plink.

Biological pathway-based associations were tested using the GSEA software package with gene sets from MSigDB^19^. Ranked lists of genes were created based on the p-values obtained from the SKAT tests. The pre-ranked GSEA algorithm with an unweighted scoring scheme was run against the hallmark, curated, GO, and immunologic signatures MSigDB gene sets for each analysis cohort.

Meta-analysis across the three sequencing batches was performed with the METAL software package^20^. The p-values and log odds ratios from Plink were used as input to the meta-analysis. The standard error analysis scheme was used, which weights effect size estimates using the inverse of the corresponding standard errors. Genomic control correction was also applied.

False discovery rates were estimated using simulated association analyses, which were run with randomly assigned case and control groups containing the same numbers of samples as the original data. Significant associations from these simulations should represent false positives.

### 2.5 Power analysis

Fisher’s exact test power calculations were made using the “statmod” R package. Power was calculated for seven control allele frequency values and eight sample sizes with a range of case allele frequency values from 0 to 1.

## 3 Results

### 3.1 Clinical characteristics

The clinical characteristics of the 118 CAN and 112 TX patients are shown in **Table 1**. The only significant variables between the patient groups were a higher percentage of Hispanic donors (p=0.008), higher rate of Type 1 diabetes in the CAN groups and an expectedly higher creatinine in CAN patients p=0.05.

**Table 1.**
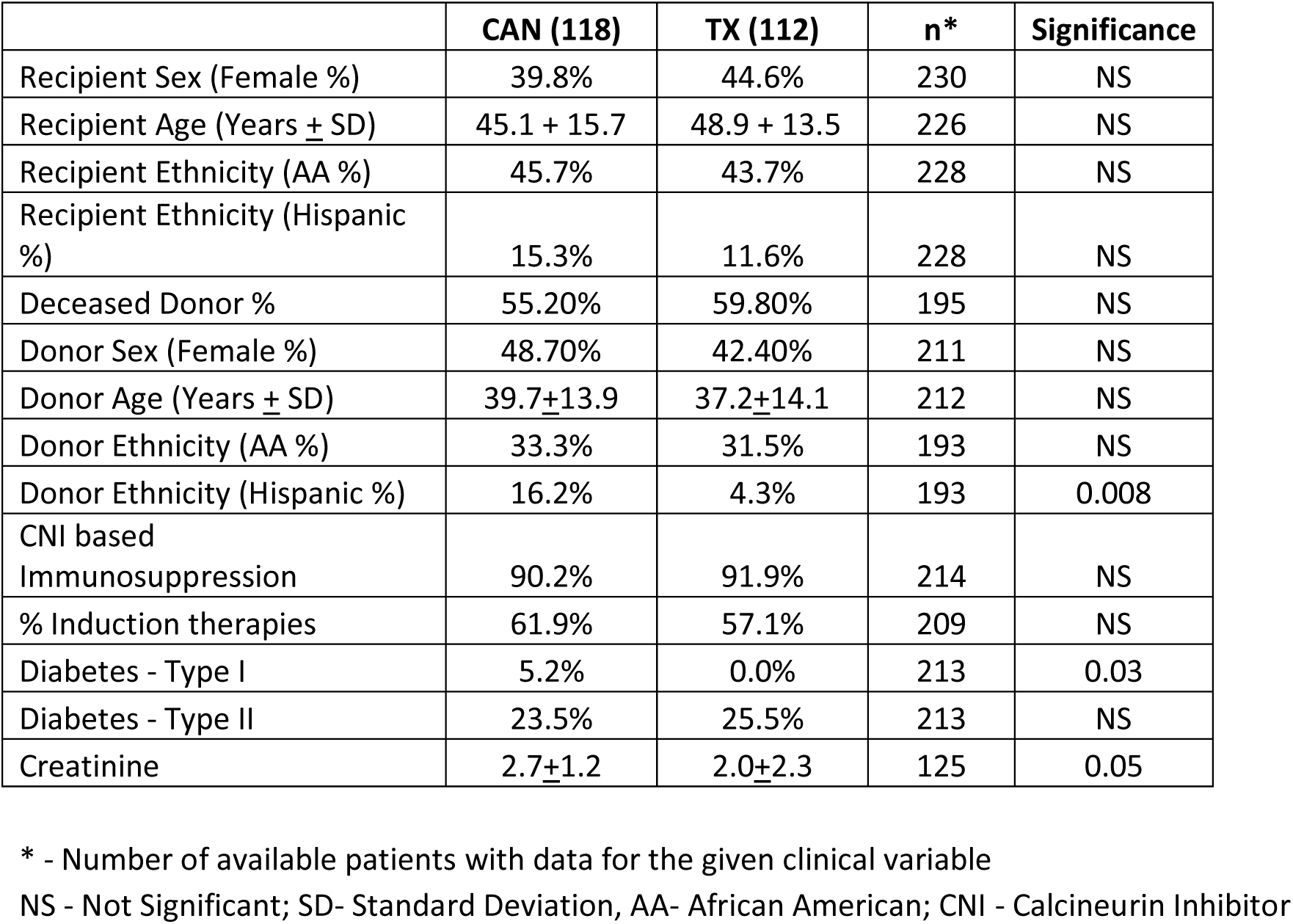
Patient demographics

|  | **CAN (118)** | **TX (112)** | **n\*** | **Significance** |
| --- | --- | --- | --- | --- |
| Recipient Sex (Female %) | 39.8% | 44.6% | 230 | NS |
| Recipient Age (Years ± SD) | 45.1 + 15.7 | 48.9 + 13.5 | 226 | NS |
| Recipient Ethnicity (AA %) | 45.7% | 43.7% | 228 | NS |
| Recipient Ethnicity (Hispanic %) | 15.3% | 11.6% | 228 | NS |
| Deceased Donor % | 55.20% | 59.80% | 195 | NS |
| Donor Sex (Female %) | 48.70% | 42.40% | 211 | NS |
| Donor Age (Years ± SD) | 39.7±13.9 | 37.2±14.1 | 212 | NS |
| Donor Ethnicity (AA %) | 33.3% | 31.5% | 193 | NS |
| Donor Ethnicity (Hispanic %) | 16.2% | 4.3% | 193 | 0.008 |
| CNI based Immunosuppression | 90.2% | 91.9% | 214 | NS |
| % Induction therapies | 61.9% | 57.1% | 209 | NS |
| Diabetes - Type I | 5.2% | 0.0% | 213 | 0.03 |
| Diabetes - Type II | 23.5% | 25.5% | 213 | NS |
| Creatinine | 2.7±1.2 | 2.0±2.3 | 125 | 0.05 |

\* - Number of available patients with data for the given clinical variable
NS - Not Significant; SD- Standard Deviation, AA- African American; CNI - Calcineurin Inhibitor

### 3.2 Sequencing and variant calling

In total, high quality exomes from 230 patients were sequenced in three batches (**Figure 1**). Patients with excellent transplant outcomes and patients who displayed extreme CAN/IFTA were roughly equally represented (112 and 118, respectively). Self-reported “white” and “black” individuals accounted for the majority of patients (86 and 103, respectively), while individuals of self-reported ancestry other than “white” and “black” made up the remaining 41 samples.

**Figure 1.**
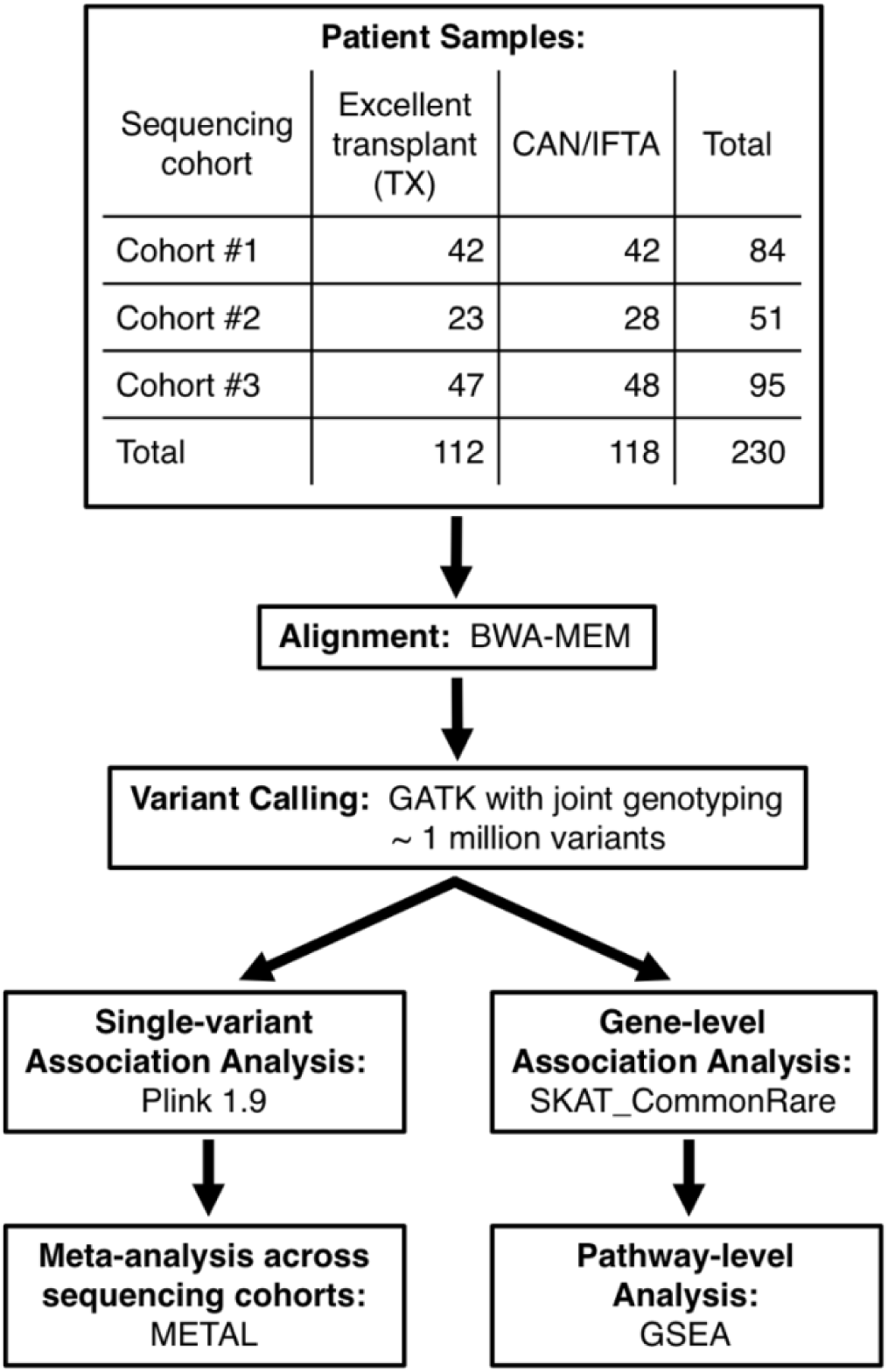
Analysis workflow and utilized software. After sample collection, sequencing, and variant calling, multiple association tests were performed. For the tests of association for individual variants, fisher’s exact tests were performed with Plink for each of the three sequencing cohorts. A meta-analysis with METAL was then performed across these three batches of sequencing. For the association tests for groups of variants, SKAT was used to rank genes by cumulative variant impact. This ranked list of genes was then used for a gene set enrichment analysis with GSEA.

On average, around 50 million (99%) read pairs from each sample were mapped to the hg19 human reference genome, and roughly 40 million (80%) of these read pairs were mapped to the whole-exome capture kit regions. The average depth of coverage across the targeted capture regions for all samples was greater than 80x, which gave sufficient power to detect single-nucleotide variants with the GATK Haplotype Caller.

The GATK Haplotype Caller identified over a million variants across all samples. Compared to the reference genome, ~470 thousand single-nucleotide and small insertion/deletion variants were identified in the first sequencing cohort, ~450 thousand variants were identified in the second sequencing cohort, and ~880 thousand variants were identified in the third sequencing cohort.

### 3.3 Association analyses

The analysis workflow for testing the association of variants with the CAN/IFTA phenotype is provided in **figure 1**. Association was first investigated at the single-variant level using the Plink 1.9-implemented Fisher’s exact tests with Lancaster’s mid-p adjustment and Monte Carlo permutations. Quantile-quantile and Manhattan plots were generated for each single-variant association analysis to visualize the distribution of obtained p-values. The full array of visualized results is provided in the supporting information.

No single variant was associated with the CAN/IFTA phenotype at a genome-wide significance level of 5×10-7. Furthermore, while they can be more sensitive than Bonferroni-corrected significance levels, simulation-based estimates of false discovery rates failed to uncover significant associations for any single variant. Tables of the top p-values from random shuffling of case and control groups are provided in the supporting information.

Given both the complex etiology of CAN/IFTA rejection and the lack of significantly associated single-nucleotide variants, we hypothesized that association tests at the gene and/or pathway level might be better able to detect a signal from aggregated single-nucleotide variants than from individual variants. However, sequence kernel association tests (SKAT) using all identified single-nucleotide variants failed to detect significant associations that either stood out from the other results or replicated across analysis cohorts. Likewise, pathway-level association tests based on gene set enrichment analysis (GSEA) of the SKAT-ranked gene associations did not produce significant results. The tables of ranked genes and pathways from these analyses are provided in the supporting information.

The final association analysis that we performed was a meta-analysis across the three sequencing batches. The top three variants that are positively associated with CAN/IFTA are displayed in a Manhattan plot in **figure 2**. This analysis also failed to pass either genome-wide significance or false-discovery tests (supporting information). However, to aid future studies, we have made the results from each of our association tests publicly available.

**Figure 2.**
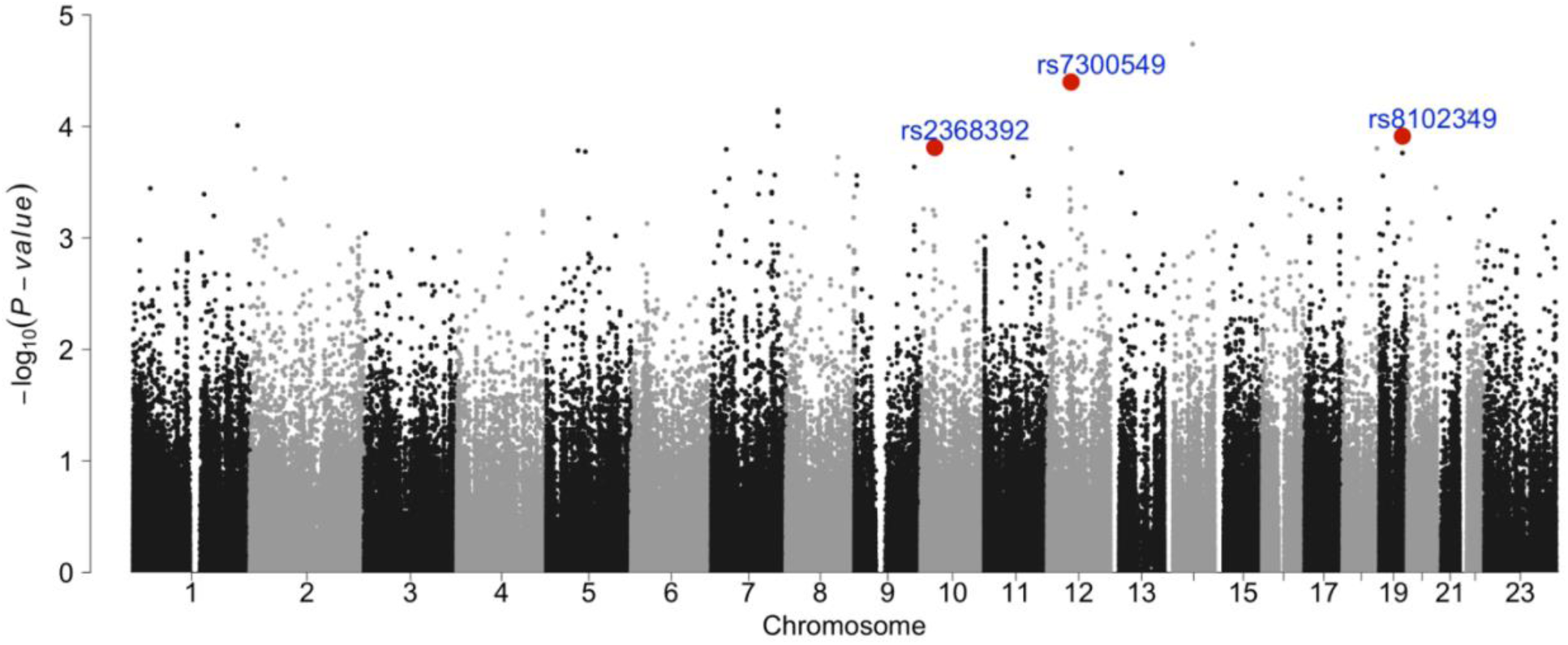
Manhattan plot of meta-analysis results. The p-values of all variants are plotted for the whole exome. The top three variants from the meta-analysis that showed positive association with CAN/IFTA for all three sequencing cohorts are highlighted.

To give some insight into our top associations, we have listed the details of the top three variants that are positively associated with CAN/IFTA in **figure 3**. The most significantly associated variant from the meta-analysis was rs7300549, with a p-value of 3.99e-05 and log odds ratio of 0.4423. This SNP is located in the LIMA1 (LIM domain and actin binding 1) gene, also known as EPLIN (epithelial protein lost in neoplasm).

**Figure 3.**
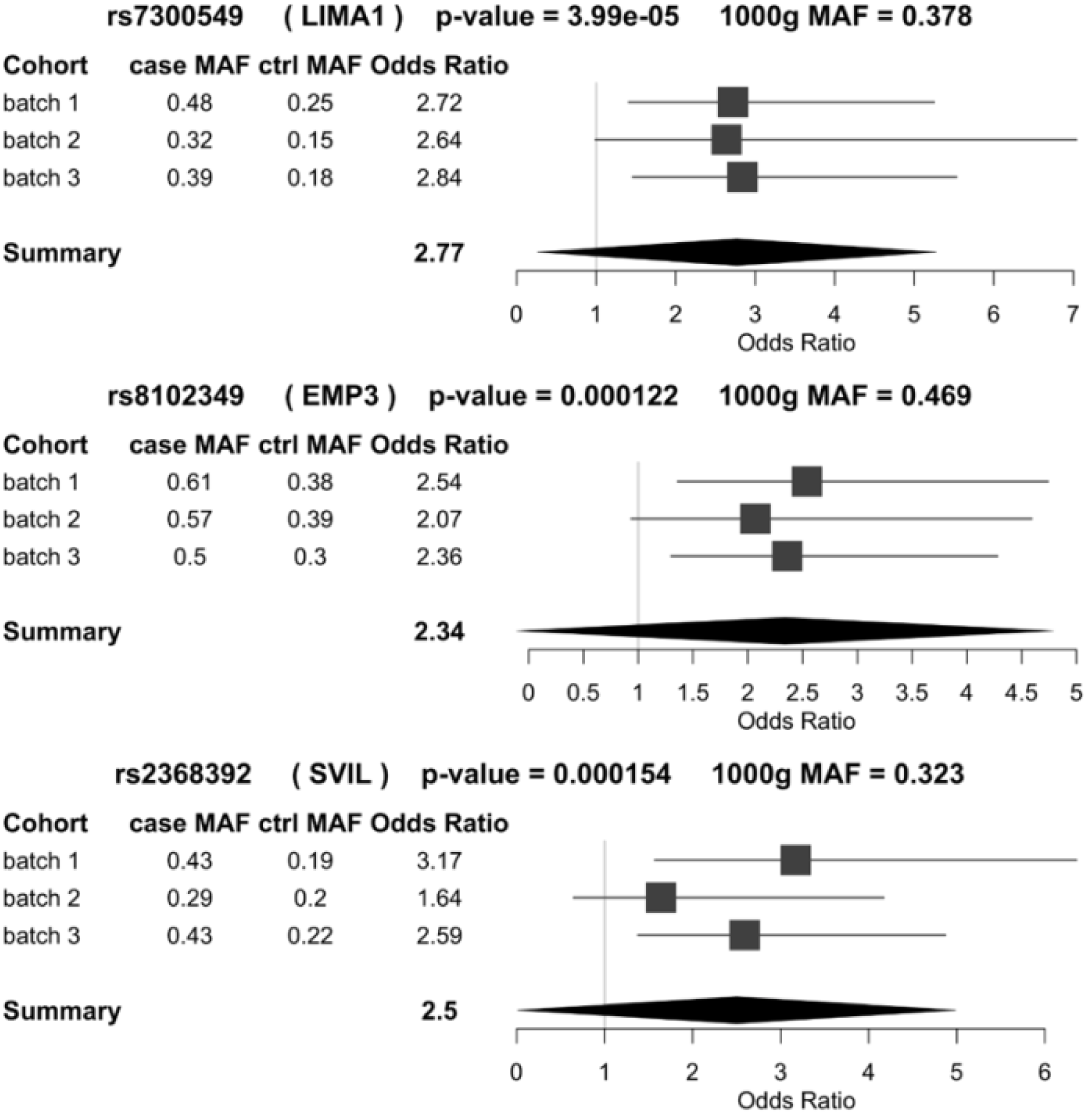
Forest plot of the three positively-associated variants with the lowest p-values. The variant rsID, gene, p-value, and 1000 genomes minor allele frequency (MAF) are above each plot. The case and control (ctrl) minor allele frequencies for the three sequencing cohorts, along with the odds ratio from the fisher’s exact test within each cohort are in the table. The odds ratio with upper and lower 95% confidence intervals are plotted for each sequencing cohort and the summary statistics for the association test are in the bottom row of each plot.

The second most significantly associated variant was rs8102349, an intron variant in the EMP3 (epithelial membrane protein) gene. EMP3 is thought to be involved in cell proliferation and cell-cell interactions and to function as a tumor suppressor^21,22^. The third most significantly associated variant was rs2368392. This SNP lies in the SVIL (supervillin) gene, which is a link between the actin cytoskeleton and the plasma membrane and plays a role in cell motility^23^.

## 4 Discussion

With a sample size of less than 250 patients, these association analyses were sufficiently powered to detect associations for common variants with a relatively large effect on the CAN/IFTA phenotype (supporting information). No significant associations were detected in our analyses, but we cannot rule out small effects from common variants or even substantial effects from relatively rare variants. On the other hand, small effects and rare variants are expected to be of little clinical utility, meaning that a low likelihood of common variants of large effect size is a useful observation, and perhaps, clinicians should not be too concerned with underlying genetic variation when treating kidney transplant recipients.

A lack of significant genetic associations with long-term renal allograft failure has been observed in previous studies. A recent study from Hernandez *et al*. looked at an order of magnitude more samples but failed to find associations outside of the HLA region^24^. The possibility that rare variants contribute to the prolonged function of kidney transplants cannot be ruled out at this time. Future studies will need much larger sample sizes to have a chance at teasing out these effects.

Despite only identifying a few associations that neared genome-wide significance, we would like to discuss the top three genes from our results in the context of the association of these genes with CAN, based on available literature. LIMA1 is an actin-associated molecule that plays a role in the development and progression of various cancers^25^. Podocytes, which are terminally differentiated cells of the glomerulus, have actin-based membrane extensions called foot processes. Kidney function has been associated with genetic mutations in proteins implicated in maintaining podocyte integrity, such as inverted formin-2 (INF2) nephrin (NPHS1), CD2-associated protein (CD2AP), α-actinin 4 (ACTN4)^26–29^. Interestingly, another LIM domain-containing protein with a role in actin polymerization, LIMK2, was shown to be associated with nephropathy in a genome-wide association study of African American diabetic kidney disease^30^.

EMP3 is a downstream effector of TWIST1/2 and induces epithelial-to-mesenchymal transition (EMT) in gastric cancer^31^. Therefore, it is tempting to speculate a major role for this gene in the EMT mechanism which has been widely implicated as a key mechanism by which injured renal tubular cells transition to mesenchymal cells leading to the development of fibrosis, a hallmark of CAN.

Like the other genes mentioned above, supervillin is also associated with the EMT process. A recent study showed that EMT due to hypoxia in hepatocellular carcinoma induces an up regulation of SVIL, which promoted cancer cell migration and invasion. This increase was a significant and independent predictor of cancer metastasis.

While our study failed to detect genetic associations, we hope that our data will be put to use in future large-scale studies and meta-analyses. The use of direct whole-exome sequencing provides a more complete set of genomic variants than commonly used microarray technology, which should allow our data to be easily incorporated into future data sets and reduce the reliance on imputation. In addition to our raw sequencing data, we have also made our full sets of both analysis code and results available for future use to ensure that our work can be efficiently integrated into larger studies.

Our sequencing data and variant calls are available at dbGaP (submission in process). The analysis results, including the results of each association test, are available on Zenodo (https://zenodo.org/record/1453460). The code to reproduce these results is on github at https://github.com/SuLab/kidneyMetaAS and can be reused and hacked for any purposes.

## Acknowledgements

This work was supported by grants awarded by the National Institutes of Health to D.R.S. (U19 AI063603) and by a TL1 award (TL1 TR002551) to L.G. through the Scripps Research Translational Institute (UL1 TR002550).

## Abbreviations

CAN: chronic allograft nephropathy
IFTA: interstitial fibrosis and tubular atrophy

## Author contributions

D.R.S. conceived the study and procured the samples. L.G. conceptualized and performed the analyses and drafted the manuscript. S.K. and T.S.M. managed patient phenotype data. L.B. coordinated the transfer of sequencing data. P.Y.K. supervised the sequencing and suggested revisions to the manuscript. A.I.S. supervised the data analysis and suggested revisions to the manuscript. All authors have read and approved the manuscript.

## Supporting information

Additional Supporting Information may be found online in the supporting information tab for this article. Quantile—quantile plots and Manhattan plots for the three sequencing cohorts and the final meta-analysis are provided in figures S1—S8. Plots of power for varying sample sizes and control group allele frequencies are provided in figures S9—S15.

